# Choreography of budding yeast chromosomes during the cell cycle

**DOI:** 10.1101/096826

**Authors:** Luciana Lazar-Stefanita, Vittore F. Scolari, Guillaume Mercy, Agnès Thierry, Héloise Muller, Julien Mozziconacci, Romain Koszul

## Abstract

To ensure the proper transmission of the genetic information, DNA molecules must be faithfully duplicated and segregated. These processes involve dynamic modifications of chromosomes internal structure to promote their individualization, as well as their global repositioning into daughter cells (Guacci et al., 1994; Kleckner et al., 2014; Mizuguchi et al., 2014). In eukaryotes, these events are regulated by conserved architectural proteins, such as structural maintenance of chromosomes (SMC *i.e.* cohesin and condensin) complexes (Aragon et al., 2013a; Uhlmann, 2016). Although the roles of these factors have been actively investigated, the genome-wide chromosomal architecture and dynamics both at small and large-scales during cell division remains elusive. Here we report a comprehensive Hi-C (Dekker et al., 2002; Lieberman-Aiden et al., 2009) analysis of the dynamic changes of chromosomes structure over the *Saccharomyces cerevisiae* cell cycle. We uncover specific SMC-dependent structural transitions between the different phases of the mitotic cycle. During replication, cohesion establishment promotes the increase of long-range intra-chromosomal contacts. This process correlates with the individualization of chromosomes, which culminates at metaphase. Mitotic chromosomes are then abruptly reorganized in anaphase by the mechanical forces exerted by the mitotic spindle on the centromere cluster. The formation of a condensin-dependent loop, that bridges centromere cluster with the cenproximal flanking region of the rDNA, suggests that these forces may directly facilitate nucleolus segregation. This work provides a comprehensive overview of chromosome dynamics during the cell cycle of a unicellular eukaryote that recapitulates and unveils new features of highly conserved stages of the cell division.

## Introduction

The improper coordination of chromosomes condensation and segregation can lead to important structural abnormalities, and result in cell death or diseases such as cancer. Pioneer studies on yeasts proved essential for the identification of genes involved in these processes. Mutations in cell-division cycle (*cdc*) (Hartwell et al., 1973) genes can block the cell cycle progression, enabling the study of global and/or local chromosome reorganization at specific cycle phases (Guacci et al., 1994; Hartwell et al., 1973; Renshaw et al., 2010; Rock and Amon, 2011; Sullivan et al., 2004; Yu, 2002). The evolutionary conserved SMC proteins bind to chromosomes in spatially and temporarily regulated manners along the cycle (Aragon et al., 2013b; Renshaw et al., 2010; Uhlmann, 2016). Cohesins, such as yeast Scc1, promote sisterchromatid cohesion during DNA replication (Blat and Kleckner, 1999; Glynn et al., 2004; Laloraya et al., 2000) and get cleaved in anaphase (Stephens et al., 2011; Uhlmann et al., 1999). At this stage, condensins such as yeast Smc2 are loaded onto sister-chromatids, facilitating their segregation (Guacci et al., 1994; Hirano, 2012; Renshaw et al., 2010; Stephens et al., 2011). A recent chromosome conformation capture study on mammalian cells has provided important insights on the organization of mitotic chromosomes’ internal structure (Naumova et al., 2013). However, no comprehensive analysis of the 4D dynamics of the chromosomes during an entire cell cycle has been achieved. To recapitulate and explore new features of cell cycle progression, we analyzed the internal folding and overall organization of *S. cerevisiae* eu- and heterochromatin over 20 synchronized time-points using chromosome conformation capture (Hi-C) (Dekker et al., 2002; Lieberman-Aiden et al., 2009).

## Results and discussion

### Cell cycle synchronizations

Hi-C libraries were generated from cell cultures synchronized in G1 with elutriation (Marbouty et al., 2014) and/or arrested at different stages of the cell cycle through thermosensitive (ts) *cdc* mutations (Fig. 1a) (Hartwell et al., 1973), sequenced, and the corresponding normalized genome-wide contact maps generated (5kb bins; Fig. 1c-d left panels; Supplementary Fig. 1; Materials and Methods) (Cournac et al., 2012). 2D maps were translated into 3D representations to visualize the main folding features (Lesne et al., 2014) (*i.e.* the centromeres and telomeres clustering in G1, Fig. 1b). Differences between two conditions were identified from log-ratio maps (50kb bins; Materials and Methods), which reflect local variations in contact frequencies. As expected, the contact map ratio of two independent G1 cell populations (experimental replicate; Fig. 1c, left panel) displays no variations. On the other hand, the ratio between exponentially growing G1 and quiescent G0 cells contact maps (Fig. 1c, lower panel) highlights the formation of the telomeres hypercluster characteristic to this metabolic condition (Guidi et al., 2015) (Fig. 1d, black arrowheads). Multiple maps can be compared by computing their pairwise distance, showing that major changes are taking place at metaphase/anaphase transition (Fig. 1e). The overall similarities/differences between datasets is summarized using principal component analysis (PCA, Fig. 1f). While experimental duplicates (such as G1, or anaphase *cdc15*) cluster together, a progression is observed between G1 (elutriated and *cdc6*) metaphase (*cdc20*) and the distant anaphase (*cdc15*).

**Figure 1.**
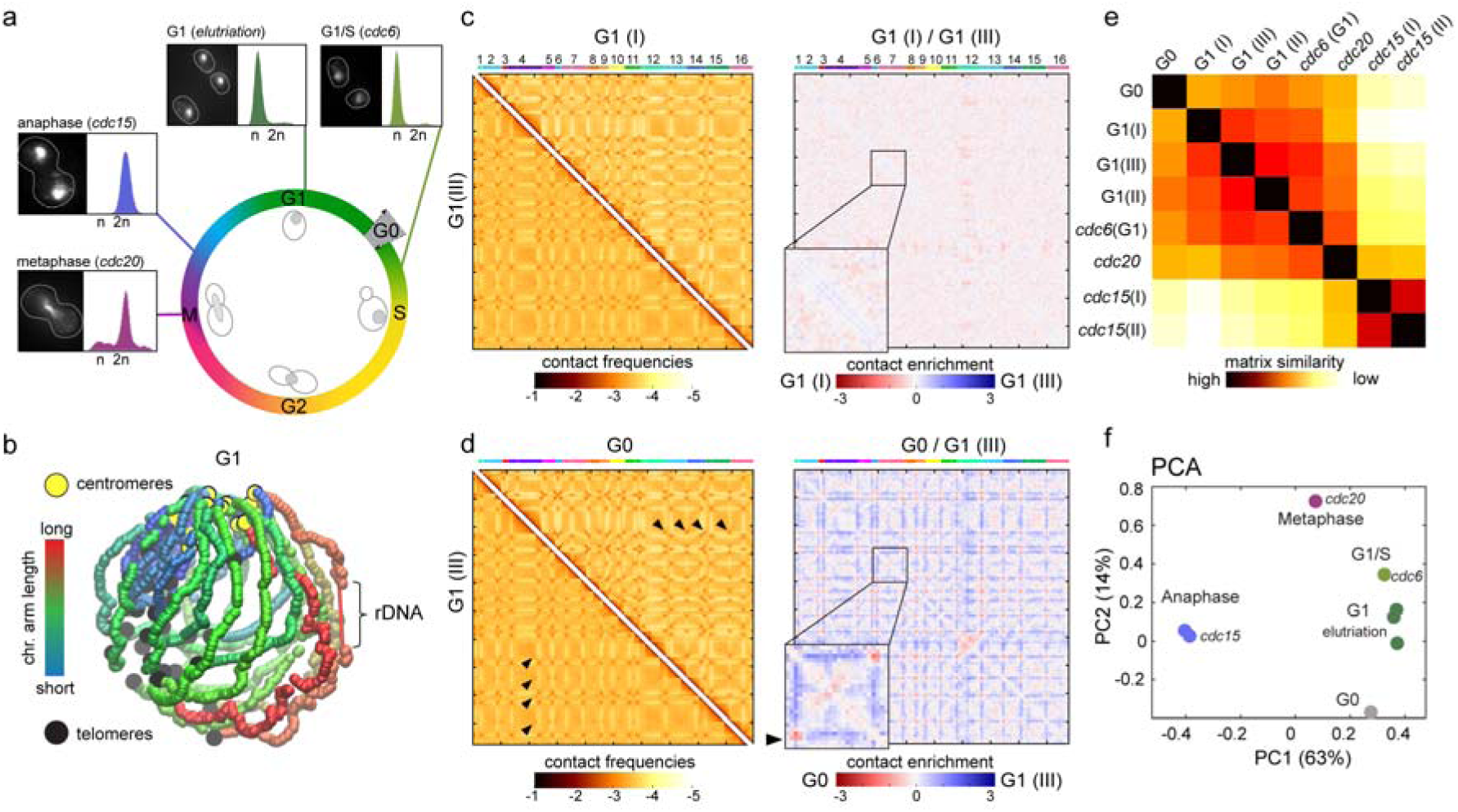
Comparison of genome structures recovered from five synchronization procedures over the cell cycle. (a) Overview of the different synchronization time-points with corresponding FACS profiles and representative images of DAPI-stained cells. (b) 3D average representation of the Hi-C contact map of a yeast G1 population. The color code reflects chromosomal arm lengths and centromeres, telomeres and rDNA are highlighted. (c-d) Contact maps comparison. The 16 yeast chromosomes are displayed atop the maps. Black arrowheads: inter-telomere contacts. Left panels display Hi-C maps obtained from (c) two G1 cell populations synchronized independently and from (d) G1 and G0 populations. Brown to yellow color scales reflect high to low contact frequencies, respectively (log10). Right panels: log-ratio between each pair of maps. The left corner boxes display magnification of chr4. Blue to red color scales reflect the enrichment in contacts in one population with respect to the other (log2). (e) Pairwise Euclidian distances between contact maps of populations of G0, G1 synchronized either with elutriation or blocked using a *cdc6* mutant, metaphase (*cdc20* mutant), and anaphase (*cdc15* mutant) cells. Color code: contact map similarity. (f) Principal Component Analysis (PCA) of the distance matrix in (e).

### Cohesins mediate chromosome compaction during S phase

To decipher the chromosome structural changes that take place during replication, synchronized G1 cells were released into S-phase and Hi-C maps generated for three time-points (two replicates) (Fig.2a; Supplementary Fig. 2; Materials and Methods). The PCA reveals a progressive structural evolution from G1 to late S/G2 phase (Fig. 2b). The dependency of the contact probability *p* on genomic distance *s*, *p*(s), reflects the internal folding of the nucleosomal fiber (Lieberman-Aiden et al., 2009; Mizuguchi et al., 2014; Naumova et al., 2013). The *p*(*s*) shows a gradual and consistent enrichment in long-range intra-chromosomal contacts (>20kb) with respect to short-range (<10kb) during replication (Fig. 2c; Supplementary Fig. 3; Materials and Methods). This change is absent when replication is impaired, although cells enter mitosis (*cdc6* ts strain (Piatti et al., 1995), Supplementary Fig. 3a). This progression stops with the completion of S-phase when it reaches the level observed in cells arrested at the G2/metaphase transition with nocodazole, a microtubule depolymerizing drug (Jacobs et al., 1988) (Fig. 2d). Interestingly, the G2/M nocodazole synchronization (Naumova et al., 2013; Sullivan et al., 2004), has a strong effect on chromosome 12 (chr12) and on the rDNA cluster structure (Supplementary Fig. 3b-c). The crossing of the p(*s*) slopes from the early/late replication time-points occurs around 10 – 20 kb (Fig. 2c, highlighted in gray), a value coherent with the *quasi*-periodic spacing between cohesin binding sites, ~11kb on average (Blat and Kleckner, 1999; Glynn et al., 2004). In agreement with the key role of cohesin in sister-chromatids folding during replication, Scc1 (Uhlmann et al., 1999) depletion in an auxin-inducible degron *scc1-aid* strain prevents the enrichment in long-range contacts in S/G2 (Fig. 2d). This result suggests that distant cohesin binding regions may be tethered together, forming chromatin loops (Guillou et al., 2010). The Scc1-dependent reorganization of the chromatin fiber supports an individualization of the sister-chromatid pairs, supported by an overall increase in intra-contacts from 63±10% to 73±4% and a decrease in inter-chromosomal contacts (Fig. 2e; see also Supplementary Fig. 8). This individualization is accompanied by an increase in centromere clustering in G2 (Fig. 2e, top right map yellow arrowheads; 2f). On the contrary, in *scc1* G2 cells intra contacts are decreased below G1 level (Fig. 2e, bottom left map), while the major binding sites for cohesin (*i.e.* centromeres) also exhibit a reduced level of contacts (Fig. 2f; Supplementary Fig. 4). These results suggest that cohesins affect the genome organization not only in G2, through the gradual establishment of sister-chromatids cohesion and chromosome individualization, but also in G1. Although yeast chromosomes are shorter than mammalian chromosomes, they similarly change their internal conformation and individualize themselves prior entering metaphase.

**Figure 2.**
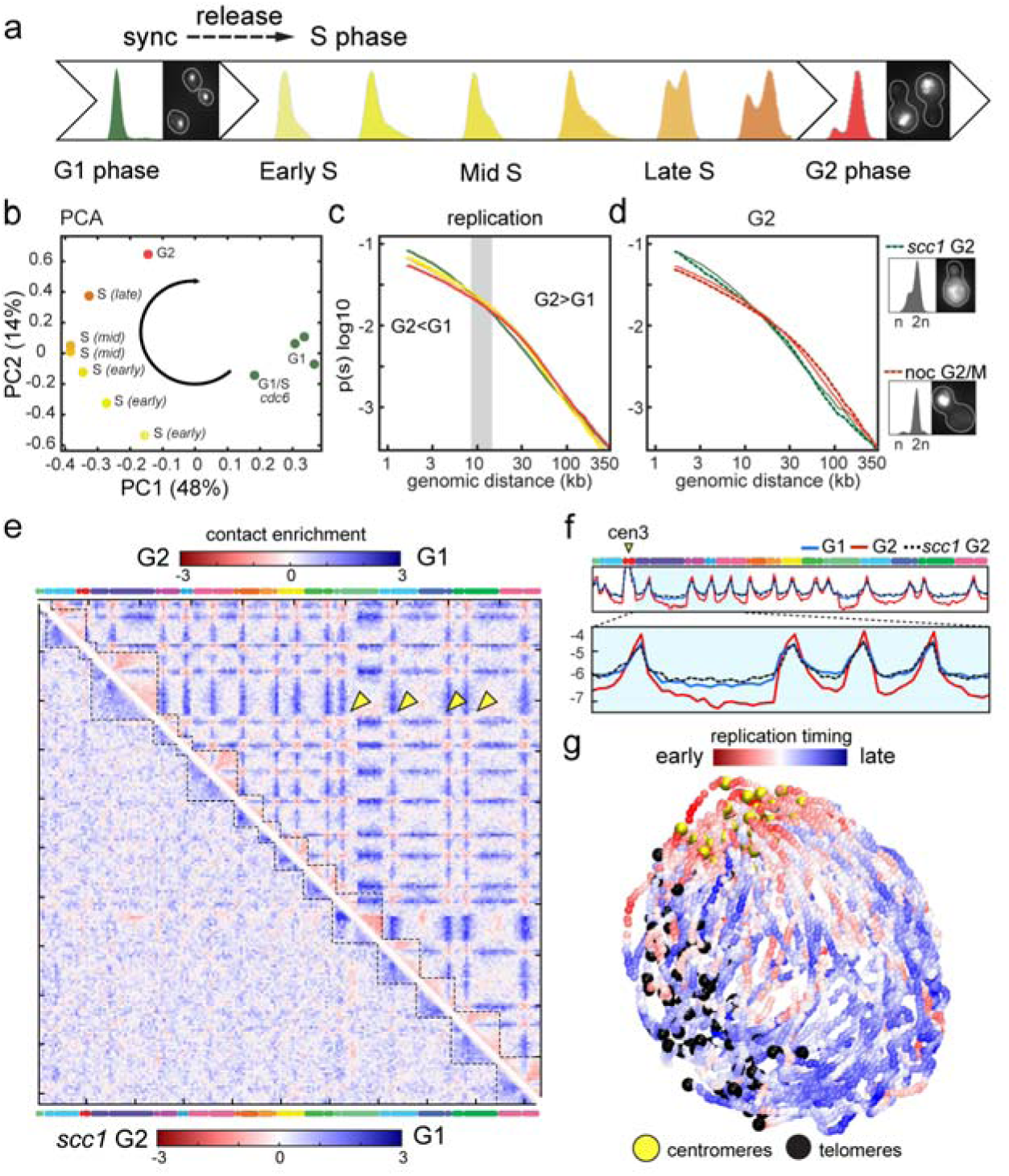
Dynamic reorganization of chromosomes during replication. (a) FACS profiles and representative DAPI-stained cells of G1 synchronized cells released in S phase.(b) PCA analysis of the distance matrix between the contact maps of the population displayed in (a).(c) p(s), *i.e.* average intra-chromosomal contact frequency (p) between two loci with respect to their genomic distance (*s*) along the chromosome (log-log scale) during replication (color code is identical to FACS profiles). (d) p(*s*) of cohesin depleted (*ssc1* G2) and nocodazole arrested (noc G2/M) cells are plotted as green and red dotted line, respectively. Corresponding FACS profiles and DAPI images are displayed on the right of the graphs. (e-f) Log-ratio of contact maps between (top right) G2 and G1 cells (yellow arrowheads: centromere contacts) and *ssc1* G2 and G1 cells (bottom left). Blue to red color scales reflect the enrichment in contacts in one population with respect to the other (log2). (f) Normalized contact frequencies between chr3 centromere (cen3) and the rest of the genome for G1, G2 and *ssc1* G2. (g) Superposition of three 3D representations of chromosomes in early replication (I, II, III). The color code indicates the replication timing. Centromeres and telomeres are highlighted.

### Spatial-temporal resolution of the replication program

In yeast, replication initiates at discrete positions. The partially stochastic and sequential activation of these replication origins defines a population-average temporal replication program (Raghuraman et al., 2001). To illustrate the link between genome organization and replication program, the sequencing coverage of the Hi-C libraries were exploited to compute the replication timing (McCune et al., 2008; Raghuraman et al., 2001) (Supplementary Fig. 5). Three early profiles were superimposed on the S-phase chromosomal structures, generating a 3D structure of the replication profile that recapitulates known properties of yeast replication and shows a “replication wave” propagating from the centromeric regions enriched in early origins onto chromosomal arms, and towards the late replicating subtelomereric regions (Fig. 2g; red and blue signal, respectively).

### Chromosome dynamics during M phase

After replication, cells progress through mitosis (M-phase), a stage regulated by multiple checkpoints. Notably, the regulatory protein Cdc20 is required for the metaphase/anaphase transition through the activation of the anaphase-promoting complex (APC) (Hartwell et al., 1973; Yu, 2002), while the Cdc15 kinase promotes mitotic exit by activating cytokinesis (Rock and Amon, 2011). Conditional mutants allow synchronization in metaphase (*cdc20*) or anaphase (*cdc15*). Contact maps of *cdc20*, *cdc15* and *cdc15*-arrested cells released into permissive conditions were generated to characterize chromosome reorganization throughout M-phase (Fig. 3a; Supplementary Fig. 6; Materials and Methods). PCA analysis shows that the major structural changes occur at metaphase/anaphase transition and that 60 minutes after release from the *cdc15* block, cells reenter a new round of replication, indicating a fully covered cycle (Fig. 3b). The p(*s*) reveals a strong increase in short-range contacts (<10-20 kb) from G2 to anaphase exceeding G1 levels and restored after anaphase completion (Fig. 3c, left panel). This increase in short-range contact is accompanied by a drop in long-range contacts, suggesting the formation of an elongated, stretched structure. Upon spindle destabilization in *cdc15*-bloked cells, the stretched chromosomal structure disappears and the two segregated chromosomal sets get closer (Fig. 3c, right panel). The latter observation is in agreement with the former report of the coalescence of the two spindle pole bodies (SPB, microtubules organizing center in yeast) in presence of nocodazole in G2/M (Jacobs et al., 1988). Altogether these results suggest that constraints resisting segregation forces remain present in anaphase, possibly resulting from the cohesion of the chromosome arm extremities. The comparison of *cdc15* and *cdc20* maps shows an increase in centromere clustering in *cdc15*, leading to the formation of a prominent polymer brush structure (Daoud and Cotton, 1982) (Fig. 3d, bottom left map, yellow arrowheads). Surprisingly, a chromosomal loop appears on chromosome 12 in *cdc15* arrested cells, bridging the centromere and the cen-proximal rDNA left flanking region (Fig. 3d box, pink arrowhead; 3e). Later in anaphase, the tel-proximal region of chr12R becomes completely isolated from the rest of the genome (Fig. 3d, top right map). 3D representations illustrate these dramatic reorganizations of the chr12 rDNA locus (3f, pink arrowheads). These results complement imaging studies showing that the rDNA exhibits a dense, line-like shape that extends throughout the nucleus at anaphase (2.1 s.d. 0.2 μm) (Sullivan et al., 2004). Upon completion of mitosis and reentry in interphase (*cdc15*+60 min), the loop disappears (Fig. 3e). Interestingly, this loop can be seen in asynchronous populations while it is only present in anaphase (Fig. 3f, lower panel).

**Figure 3.**
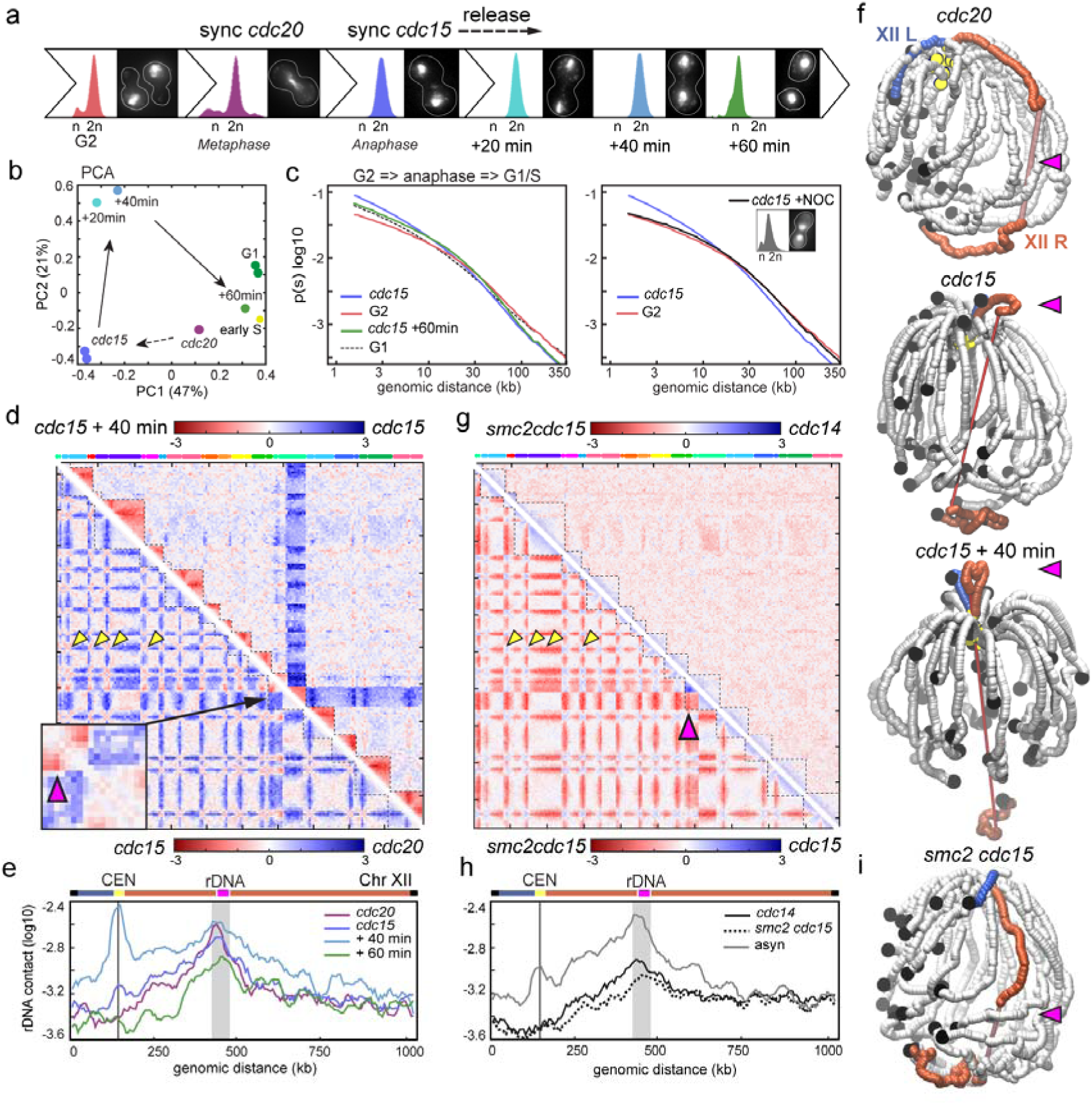
Dynamic reorganization of chromosomes during segregation. (a) FACS profiles and representative DAPI-stained cells of synchronized and/or released populations, from G2 until reentry in G1/S. (b) PCA analysis of the distance matrix between the contact maps of the populations described in (a). (c) Left panel: p(*s*) of cells in G1, G2, anaphase (*cdc15*) and released from a *cdc15* block (60 minutes). Right panel: p(*s*) of G2, and *cdc15* blocked cells (anaphase) in absence or presence of nocodazole (*cdc15*+NOC). (d) Log-ratio of contact maps between metaphase(*cdc20*) and anaphase (*cdc15*) arrested cells (bottom left) and *cdc15* and anaphase released (*cdc15* + 40min) cells (top right). Yellow arrowheads: inter-centromere contacts. Box: magnification of the contacts between the rDNA flanking region and chr12 centromere (pink arrowhead). (e) Distribution of intrachromosomal contacts of a cen-proximal rDNA flanking region (highlighted in grey) with the rest of chr12 in *cdc20*, *cdc15*, and *cdc15* release (black line: centromere). (f) 3D representations of the contact maps from *cdc20*, *cdc15* arrested and *cdc15* released cells. The right (XIIR) and left (XIIL) arms of chr12 are highlighted in red and blue, respectively. The pink arrowhead points at the right arm anaphase loop. Centromeres, telomeres and rDNA are highlighted. (g) Log-ratio maps of cells blocked in anaphase with our without condensin (*cdc15* vs. *smc2 cdc15* strain; bottom left) and of *cdc14* and *smc2 cdc15* cells (top right). (h) Distribution of intra-chromosomal contacts of a cen-proximal rDNA flanking region (highlighted in grey) with the rest of chr12 in *smc2 cdc15*, *cdc14* and asynchronous contact map. (i) 3D representation of the contact map from smc2 *cdc15.*

### Condensation is a locus specific process that occurs in anaphase

The proper condensation and segregation of the rDNA cluster requires the Smc2 condensin and the nucleolar release of the Cdc14 phosphatase (Clemente-Blanco et al., 2009; D’Amours et al., 2004; Sullivan et al., 2004; Yoshida et al., 2002). We investigated the influence of both factors on rDNA organization during anaphase (Supplementary Fig. 7). Smc2 depletion in *smc2-aid cdc15* strain reduces centromere clustering in anaphase and suppresses the loop, as well as chr12R extremity isolation (Fig. 3g, bottom left map; 3i). Strikingly, *smc2 cdc15* and *cdc14* maps are similar, and the loss of centromere clustering and of the chr12 anaphase loop in both mutants reveal their epistatic relationship (Fig. 3g, upper right map, Fig. 3h). Since both mutants exhibit difficulties to complete anaphase, the condensin-dependent loop may play a direct role in the segregation of the rDNA cluster through the application of a pulling force directly onto the rDNA region.

## Conclusions

A global pattern of structural changes during the cell cycle can be summarized from centromere contacts, intra/inter contact ratio and short/long contact ratio computed for each of the 20 time-points(Fig. 4). First, short/long contact ratio recapitulates the three internal folding states of chromosomes (G1, G2, and anaphase; Fig. 4; Supplementary Fig. 8). Second, intra/inter contact variations reflect the successive phases of chromosome individualization and intermingling. Individualization occurs at the end of replication and peaks during anaphase resolution. Finally, the intra/inter ratio correlates strongly with centromere clustering (c=0.72, p=10^−4^), with both ratios peaking during anaphase exit. A possibility is that the strengthening of polymer brush organization contributes to chromosome individualization. Overall, this comprehensive analysis of the 3D chromosome choreography during replication and segregation brings to light new perspectives regarding these fundamental processes.

**Figure 4.**
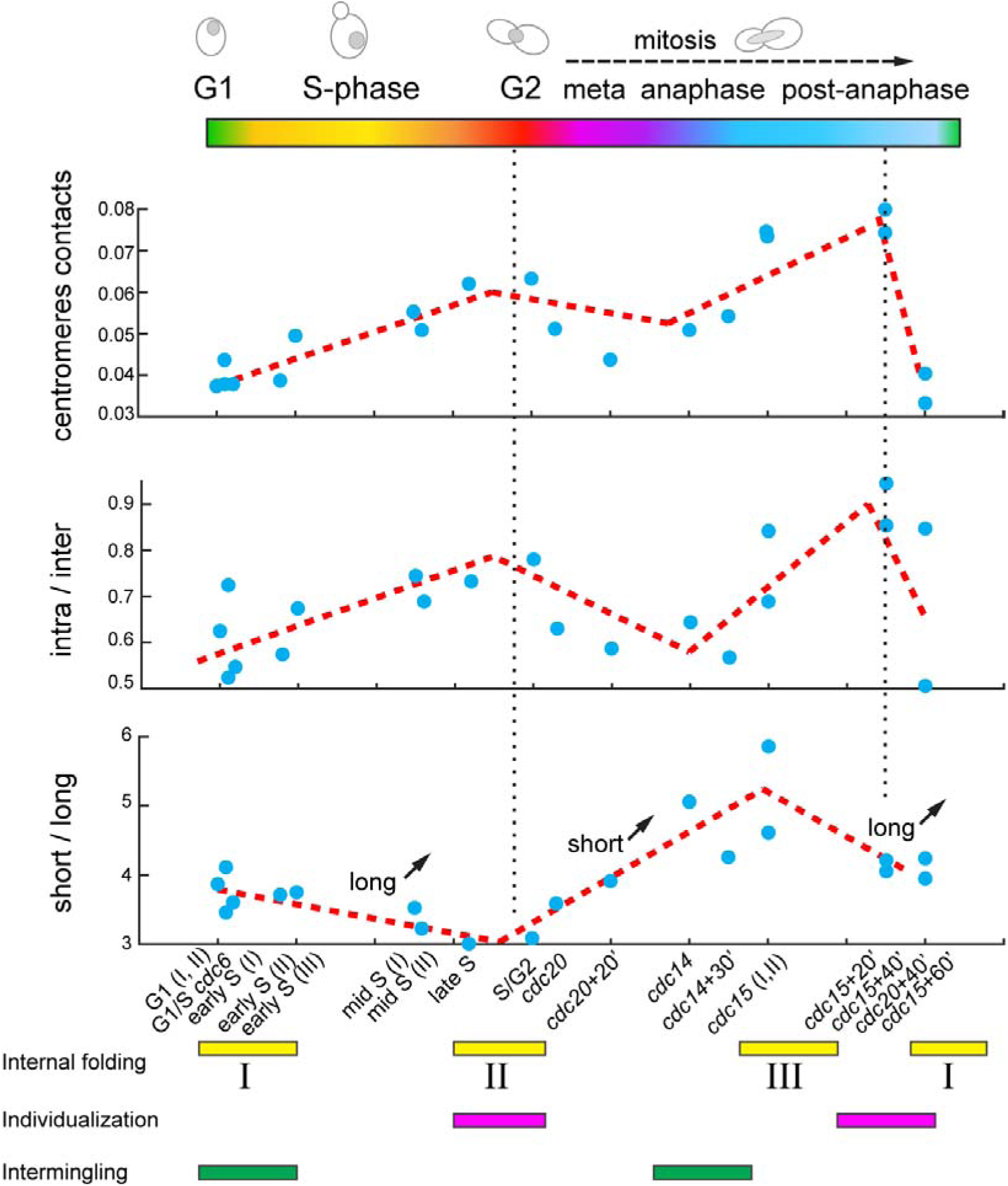
Spatio-temporal reorganization of yeast genome. Dynamics of centromere contacts, intra/inter contact ratio and short/long contact ratio for each of the 20 time-points (blue dots), covering the entire cell cycle (upper panel). Correlation with the three chromatin folding states (bottom panel).

## Sample preparation

### Media and culture conditions

Yeast strains used in this work are listed in Supplementary Fig. 9. All strains were grown in rich medium (YPD: 1% bacto peptone (Difco), 1% bacto yeast extract (Difco) and 2% glucose), except for YKL051 (*MET3-HA-CDC20*) that was grown in synthetic complete medium deprived of methionine (SC: 0.67% yeast nitrogen base without amino acids (Difco), supplemented with a mix of amino-acids, uracil and adenine, 2% glucose). Cells were inoculated and grown overnight in 15 ml of growth medium at either 30°C or 23-25°C (the later temperature corresponding to the permissive temperature of the conditional temperature-sensitive mutations *cdc6-1*, *cdc14-3* and *cdc15-2*). The next morning 2 ×10^9^ cells of BY4741 and YKL050 strains were restarted in 150 ml fresh YPD at 30°C for 2 h and fixed for Hi-C as asynchronous state. On the contrary, to obtain synchronized cell populations, 10 ml of the overnight cultures were diluted into 800 ml of the corresponding growth medium (YPD or SC) and grown overnight at 30°C or 23-25°C. Cells were then harvested through centrifugation, washed and suspended in 1X PBS, and elutriated using Beckman Avanti J-26 XP elutriation system (JE-5.0 elutriator rotor). Fractions of approximately 1-3 × 10^9^ G1 daughter cells were collected and eventually restarted in the adequate medium and temperature conditions so as to obtain synchronized fractions that are well-distributed across the cell cycle (see further synchronization details). On the other hand, the dataset corresponding to the quiescent state (G0) is coming from already published data in (Guidi et al., 2015), and was obtained by carbon source exhaustion.

### Synchronization with elutriation (G1 cells)

To recoverG1 daughter cells, the exponential growing cultures were elutriated - a physical method of synchronization, used to separate cells according to their density and sedimentation velocity (Marbouty et al., 2014). Briefly, the 800 ml overnight culture was centrifuged, washed in 1X PBS and pelleted cells were suspended in 1000 ml of fresh YPD for 2 h at 30°C. This additional growing step allowed cells in stationary phase to reenter exponential phase before being elutriated. For each elutriation experiment, 2-1.8 × 10^11^ cells were washed and suspended in 30 ml of 1X PBS and injected in the 40 ml elutriation chamber at an average flow rate ranging from 20 ml/min to 25 ml/min (MasterFlex L/S pump from Cole-Parmer), at 2500 r.p.m. and 23°C. Cells were then left to equilibrate in 1X PBS for 45 min at a constant flow and rotational speed. To start collecting the first fractions containing the small G1 cells, a periodic 2 ml/min increment of the flux was applied between each fraction. The resulting 600 ml fractions were centrifuged and approximately 2.5 × 10^9^ G1 cells/fraction were recovered. Before fixating the G1 state, cells were suspended in fresh YPD at 30°C for 30 min, so they could recover from their stay in PBS during the elutriation. To minimize the potential variability introduced by the age heterogeneity of the bulk population, G1 daughter cells were used as starting point for all cell cycle synchrony and in combination with genetic and chemical synchronization methods (see below).

### Release into S phase

To analyze cells undergoing replication, G1 elutriated cells were released into S phase. 2 ×10^9^ overnight G1 cells – belonging to the same elutriated fraction to minimize S phase restart heterogeneity – were inoculated into 150 ml YPD at 25°C (to lower the speed of the replication forks). Upon release, aliquots were sampled all over S phase till G2 with an average periodicity of 5 min. Apposite kinetics, meant to monitor the restart of the overnight G1 cells, allowed us to estimate an approximate time lag of 2h10min at 25°C required for the S phase to begin. Therefore, the restarted aliquots were fixated for Hi-C at: 2h15min, 2h20min, 2h25min, 2h30min, 2h35min, 2h40min and 2h45min. The progression of each fraction throughout the S phase (from G1 to G2) was monitored with flow cytometry.

### Synchronization through thermosensitive mutations

The genetic synchronizations, using thermosensitive (ts) *cdc* strains (Hartwell et al., 1973), were all performed starting from elutriated G1 daughter cells, released in temperature conditions designed to arrest the progression of the cycle at specific phases. The G1/S checkpoint synchronization was obtained using the YKL054 strain (caring the *cdc6-1* mutation). The YKL054 was grown overnight at 25°C. Cells were then restarted in fresh media the next morning, and elutriated while in exponentially growing stage. The elutriated G1 cells were incubated in fresh YPD at the non-permissive temperature of 37°C for 3 h. The same procedure was employed to synchronize cells during M phase, - in early (strain YKL052, *cdc14-3*) and late anaphase (strain YKL053, *cdc15-2*) states, respectively. Therefore, G1 cells of both YKL052 and YKL053 ts strains were incubated for 3 h at the non-permissive temperatures of 30°C and 37°C before being processed for Hi-C. Moreover, to investigate the dynamics of mitotic exit and cell cycle reentry, several anaphase synchronized aliquots were shifted at the permissive temperatures of 23°C and 25°C and different time-points were taken (YKL052: 30 min; YKL053: 20 min, 40 min and 60 min). To study non-replicated mitotic chromosomes, the YKL054 strain was kept in non-permissive growing conditions for an extended period of 6 h. During this time lapse G1 cells bypassed the G1/S checkpoint and went directly into M phase without replicating their chromosomes. The synchrony of each time point (in G1/S, anaphase and release) was monitored with flow cytometry and microscopy.

### Synchronization through chemical compounds

The chemical synchronization, as the genetic one, was performed on G1 elutriated cells. Synchronization at the G2/M transition was achieved by restarting G1 cells (strain YKL050) in YPD at 30°C for 1 h, followed by the addition of nocodazole (Calbiochem) 15 μg/ml and incubation for another 2 h at 30°C. Cells arrested in G2/M with nocodazole were either crosslinked, or washed and inoculated in fresh YPD at 30°C. The washing of nocodazole allowed G2/M synchronized cells to enter M phase, which was monitored at different time intervals (20 min, 45 min, 60 min and 90 min). For M phase synchronization in metaphase, we used the YKL051 strain (*MET3-HA-CDC20*). The YKL051 G1 elutriated cells were restarted in YPD complemented with 50 μg/ml methionine for 5 h at 30°C. Cells arrested in metaphase were split in different fractions: some were fixed for metaphase, other were washed and suspended in SC medium without methionine and different time-points were taken (20 min and 40 min). To study chromosome reorganization as a function of cohesin and condensin activity, we used YKL055 and YKL056 strains in which *SCC1* and *SMC2* genes, respectively, were tagged with an auxin-inducile degron (*aid*). Therefore, the degradation of these proteins was induced when auxin (IAA) was added to the medium at a final concentration of 20 μM. At first, both YKL055 and YKL056 were grown and elutriated in absence of IAA; then the G1 cells were restarted in YPD supplemented with IAA at 30°C. The cohesin *scc1-aid* mutant was processed for Hi-C in late S/G2 (see: release into S phase); while the *smc2-aid* was arrested in late anaphase using the coexistent *cdc15-2* mutations (see: synchronization through thermosensitive mutations). The synchrony of each time point was monitored with flow cytometry and microscopy. All the resulting synchronized time points were used to build Hi-C libraries.

### Flow cytometry

About 5 × 10^6^ cells were fixed in ethanol 70% and stored at 4°C overnight. Cells were then pelleted, washed and incubated in sodium citrate 50□mM (pH□7.4) complemented with RNase A (10 □mg/ml; Roche) for 2 h at 37°C. Next, Sytox green (2 μM in sodium citrate 50 □mM; ThermoFisher) was added and cells incubated for 1 h at 4°C. Flow cytometry was performed on a MACSQuant Analyser (Miltenyi Biotec) and data was analyzed using FlowJo X 10.0.7 software (Tree Star).

### Microscopy

Fractions of cells fixed in ethanol 70% and stored at 4°C overnight were pelleted and washed 3 times for 5 min in 1X PBS. Cells were permeabilized by immersion in 0.2% Triton X-100 (Biosolve) for 5 min. To remove the triton, cells were pelleted and washed 3 times in 1X PBS. The liquid was aspirated and cells were suspended in DAPI labeling solution (2 μg/ml in 1X PBS) for 10 min at room temperature. Before imaging acquisition, the labeling solution was aspirated and the cells were washed 3 times for 5 min in 1X PBS. Cells were imaged at 350 nm excitation wavelength with Nikon fluorescence microscope (Camera Andor Neo sCMOS, software Andor IQ2 2.7.1, LED Lumencor Spectra X).

## Acknowledgements

We thank Martial Marbouty and Axel Cournac for technical help in the earlier stage of this project, and Emmanuelle Fabre, Angela Taddei, Stephane Marcand, Thomas Guérin, Philippe Pasero, and Etienne Schwob for sharing strains and for discussions. Vittore Scolari and Heloise Muller were partly supported by Pasteur-Roux-Cantarini postdoctoral fellowships. This research was supported by funding to R.K. from the European Research Council under the 7th Framework Program (FP7/2007-2013, ERC grant agreement 260822), from Agence Nationale pour la Recherche (MeioRec ANR-13-BSV6-0012-02), and from ERASynBio and Agence Nationale pour la Recherche (IESY ANR-14-SYNB-0001-03).

## Authors Contributions

LLS and RK designed research. LLS performed the experiment, with help from GM, AT and HM. VS and JM analyzed the data, with contributions from LLS. LLS, JM and RK interpreted the data and wrote the manuscript.

## Authors information

Sample description and raw contact maps are accessible on the GEO database through the following accession number: xxx. Raw sequences are accessible on SRA database through the following accession number: xxx. The authors declare no competing financial interests. Correspondence and requests for materials should be addressed to R.K., or J.M.

